# Transition State Characteristics During Cell Differentiation

**DOI:** 10.1101/264143

**Authors:** Rowan D. Brackston, Eszter Lakatos, Michael P.H. Stumpf

## Abstract

Models describing the process of stem-cell differentiation are plentiful, and may offer insights into the underlying mechanisms and experimentally observed behaviour. Waddington’s epigenetic landscape has been providing a conceptual framework for differentiation processes since its inception. It also allows, however, for detailed mathematical and quantitative analyses, as the landscape can, at least in principle, be related to mathematical models of dynamical systems. Here we focus on a set of dynamical systems features that are intimately linked to cell differentiation, by considering exemplar dynamical models that capture important aspects of stem cell differentiation dynamics. These models allow us to map the paths that cells take through gene expression space as they move from one fate to another, e.g. from a stem-cell to a more specialized cell type. Our analysis highlights the role of the transition state (TS) that separates distinct cell fates, and how the nature of the TS changes as the underlying landscape changes— change that can be induced by e.g. cellular signalling. We demonstrate that models for stem cell differentiation may be interpreted in terms of either a static or transitory landscape. For the static case the TS represents a particular transcriptional profile that all cells approach during differentiation. Alternatively, the TS may refer to the commonly observed period of heterogeneity as cells undergo stochastic transitions.

## Introduction

Cells are not inert objects. They have finite lifetimes with typically well defined origins and ends. And over the course of their lifetime – which lasts anything from minutes to many years – change in response to environmental, physiological and, potentially, developmental signals (Waddington, 1957). Some of these changes are minor, e.g. changing the expression of certain proteins in response to an environmental signal, or the activity of an enzyme as part of metabolism. Others relate to longer-term, less reversible, or more profound changes in cell state; including commitment to replication, apoptosis, or differentiation. The former set of changes can be viewed as *tactical* decisions which are made in response to (typically transient) changes in a cell’s environment (Filippi et al., 2016), whereas the latter are of more *strategic* importance for the cell and, where relevant, potentially the organisms as a whole (Hormoz et al., 2016; Moris and Arias, 2017).

In humans, a single fertilised egg cell eventually gives rise to some 35 trillion cells in the adult. How many cell types there are remains an unanswered question, but some aspects of the process by which an omni-potent stem cell differentiates into a more specialised cell are now becoming clearer. Remodelling of the gene regulatory networks—typically in response to signalling events — change the transcriptional programme of the cell, thereby leading to a concomitant change in cell phenotype/state. We will here assume for ease of argumentation, that the molecular state of a cell can reflect the true state of the cell, its *phenotype*, but stress that this may only be a poor substitute for a more direct biological or phenotypic characterization.

The popular metaphor of the *Waddington’s epigenetic landscape* has come to predominate much of the discussion about cell differentiation processes: cells are described as marbles rolling through a landscape of hills and valleys drawn towards local points of minimum elevation (Waddington, 1957). An individual ball starts its journey in a valley at the back of the landscape and as it progresses forward (the passage of time in the original formalism) and downward; it might face branching points along the path, representing the series of (typically binary) fate choices made by a developing cell. Every point a ball travels through represents a cellular state, for example a specific level of expressed RNA or protein. Although the number of possible states is theoretically infinite, the number of phenotypes observed in actual cells are often very limited. In this view, the final basins of low elevation where a high proportion of cells end up correspond to these experimentally observable, terminal cell types.

The key insight offered by the landscape is to illuminate how genetically identical cells can attain distinct phenotypes following differentiation, and furthermore how these phenotypes persist in daughter cells. While such persistence and memory effects are understood to result from the epigenetic and proteomic state of the cells, the landscape itself is widely regarded to be shaped by the underlying gene regulatory network (Moris et al., 2016; Huang et al., 2017). In this way, models describing the co-regulation and interactions between different genes may also be understood to describe the epigenetic mechanisms underlying persistent stem cell differentiation.

The landscape notation has been widely used as a qualitative way of understanding and illustrating the dynamics driving development (Pujadas and Feinberg, 2012; Feinberg et al., 2016; Teramoto et al., 2017). Moreover, even though Waddington may not have intended his landscapes to be any more than a conceptual tool, landscapes can be given quantitative meaning: exploring the behaviour of the underlying network in terms of a probabilistic landscape framework. A succession of studies have recently proposed different approximations to potential functions that can serve as mathematical descriptions of the epigenetic landscape for a cell fate regulatory network. Here the elevation of the surface reflects the probability of observing a particular state in phase space (Sasai and Wolynes, 2003; Wang et al., 2010a, 2011; Li and Wang, 2013; Zhang and Wolynes, 2014; Huang et al., 2017): states that have the highest probability locally will have lower potential and hence will act as the valley-bottoms on the landscape, surrounded by a basin of attraction, which in this picture would correspond to cells with slightly different states but exhibiting the same phenotype.

Even though the landscape depiction might be adopted to many decision making processes over the lifetime of a cell, its major uses are still in describing development and stem cell differentiation processes (Li and Wang, 2015; Yamanaka, 2009). In this context the final attractors of the landscape are differentiated cell types with well-defined patterns of robust gene expression, and differentiation occurs through transitions along the surface, while some cells might reside or return to the original basin of the pluripotent phenotype.

Mathematically, we can draw on a vast body of work to characterize the cell states defined in terms of steady-states of molecular concentrations. For the sake of clarity, we consider *X* to denote the state of the system (e.g. the whole set of mRNA and/or protein abundances), which evolves according to the stochastic differential equation

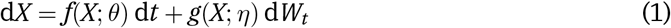

where *f*(*X*; *θ*) describes the deterministic dynamics of the system, and *g*(*X*; *η*) d*W_t_* captures the stochastic components of the dynamics. If the latter can be ignored we recover a more conventional ordinary differential equation, commonly written as

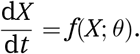

This, setting the left-hand side equal to zero allows us to solve for the (deterministically) stationary states of the system, which if they are stable, i.e. if they are *attractors*, we assume correspond to distinct cell states.

If the set {*X*_1_, *…*, *X_q_* } denotes the set of stable stationary/attractor states, we identify them with the valleys/local minima in the corresponding epigenetic landscape for the system. Classifying the stability of these solutions (e.g. in the presence of stochastic effects), and the basins of attraction has been one of the long-standing problems in dynamical systems theory. Closely linked to it is the question of how different stationary points can be reached from one another: is there a particularly favoured path that the system traverses as it moves from, say *X_i_* to *X_j_* (1 ≤*i*, *j* ≤ *q*)?

Below we will use a set of illustrative examples that allow us to study the transitions between stationary solutions of (stochastic) differential equations, with an emphasis on dynamical systems in the context of stem cell differentiation. In particular we shall be investigating the concept of *transition states* (TS), as recently discussed and reviewed by Moris et al. (2016). For representative model systems we discuss two competing definitions for the transition state in the context of static and transitory landscapes. For static landscapes we shall employ the concept of the *minimum action path* (MAP) (Freidlin et al., 2012), to examine typical transition paths. We first discuss the properties of transition states for an examplar “toy” system before applying the same analysis to a developmental model. In particular we will aim to answer four linked questions:

- What defines a TS for a stochastic dynamical system?
- How long does the system spend at the TS?
- Is the TS the same for forward and backward dynamics?
- How does the epigenetic landscape, and especially the TS change, in response to external signals?

## 1 The Potential Landscape and Transition States in a Bistable System

In order to exemplify some key properties of stochastic dynamical systems in the context of potential landscapes we provide a simple model that exhibits some of the hallmarks found in real developmental systems. The model we provide is, however, not intended to represent any particular biological process, but serves as a simple example in which potential landscapes and transition states may be described. We will use this model to firstly demonstrate the distinction between gradient and non-gradient dynamics, before examining two definitions for the concept of the transition state.

### 1.1 Limitations to gradient based dynamics

The SDE defined in equation 1 above describes a dynamical system with both deterministic and stochastic components. In general, the deterministic forcing vector *f*(*X*; *θ*) of such SDEs may be decomposed into the sum of the gradient of a potential, Δ*U*, and a remaining component often referred to as the *curl* or *flux* (Wang, 2015; Zhou and Li, 2016). This decomposition may be written as,

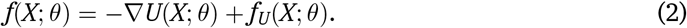

The underlying concept is that the potential describes the portion of the dynamics that determines the basins of attraction (and thus the most probable states). This potential thereby gives an intuitive explanation for the observed behaviour; the dynamical system will tend to stay near to the minima in the potential and will move away from regions of higher potential.

A number of methods exist for determining the decomposition in which the potential is defined; however, by far the most common is via the steady state probability distribution over the state space, where *U* ∝ – ln(*P_s_*(*X*)). Numerous studies have obtained landscapes based upon this approach, either using methods that directly obtain the steady state distribution (Lv et al., 2014; Ge et al., 2015; Li et al., 2016), or use extensive simulations to estimate the distribution empirically (Wang et al., 2008, 2010a; Li et al., 2011; Guo et al., 2017). Alternative approaches include those based upon variational principles and the action in moving between any two points (Wang et al., 2010b, 2011; Lv et al., 2014), empirical Lyapunov functions (Bhattacharya et al., 2011) or an orthogonal decomposition of the vector field (Zhou et al., 2012). Regardless of the method, the concept of dynamics composed of both a potential and vector field is commonplace, and the potential is indeed related to the idea of an epigenetic landscape.

In order to elucidate the distinction between gradient-based and rotational dynamics we first examine a simple bistable system expressed in SDE form. We define the system in terms of a pre-specified two-dimensional potential *U*(*X*; *θ*), chosen as,

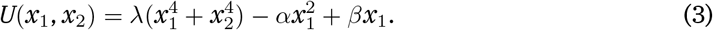

This potential is displayed in Figure 1(A),(B) and can be seen to consist of two minima at (*x*_1_, *x*_2_) ≈ (±1, 0) and one saddle point at (*x*_1_, *x*_2_) ≈ (0, 0). Given such a potential field, we may then define the vector field *f*(*X*; *θ*) as the sum of –∇*U* and a component *f_U_* chosen here to be such that ∇*U f_U_* = 0. Given the choice of *U* in equation 3, these two components are given as

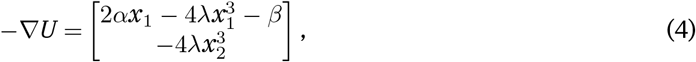

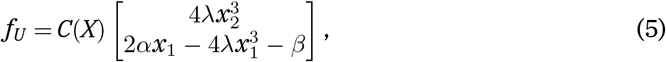

respectively, where *C*(*X*) can be any scalar state dependent function, chosen here to be a simple constant *c*. The vector field components shown in Figure 1(B) display the two components of *f*. The black vectors can be seen to point in the direction of decreasing *U* and represent the –∇*U* component. The white vectors represent the curl component of the vector field and are parallel to the contours of *U* due to the orthogonality. Together these components on average lead to spiralling paths leading towards the potential surface’s minima. With sufficient stochastic forcing the system may also make intermittent transitions between the two potential wells.

Figure 1(C) displays the probability distribution over the state space defined by the variables *x*_1_ and *x*_2_. The distribution is clearly strongly linked to the landscape displayed in Figure 1(A),(B), (it is in fact implied by it) and is seen to peak at (*x*_1_, *x*_2_) ≈ (±1, 0). By contrast the system spends relatively little time around the origin which is the third, unstable, fixed point of the system. It is worth noting that the distributions for the cases with and without a curl component are practically indistinguishable. This is typical for such systems and is the basis for identifying a landscape based upon the probability distribution alone Wang (2015). However, a corollary of this is that only part of the dynamics are readily identifiable based upon static data Weinreb et al. (2017); in a biological context, knowing the trasncriptomic states of cells tells us very little about the paths through transcription space taken during differentiation.

**Figure 1:**
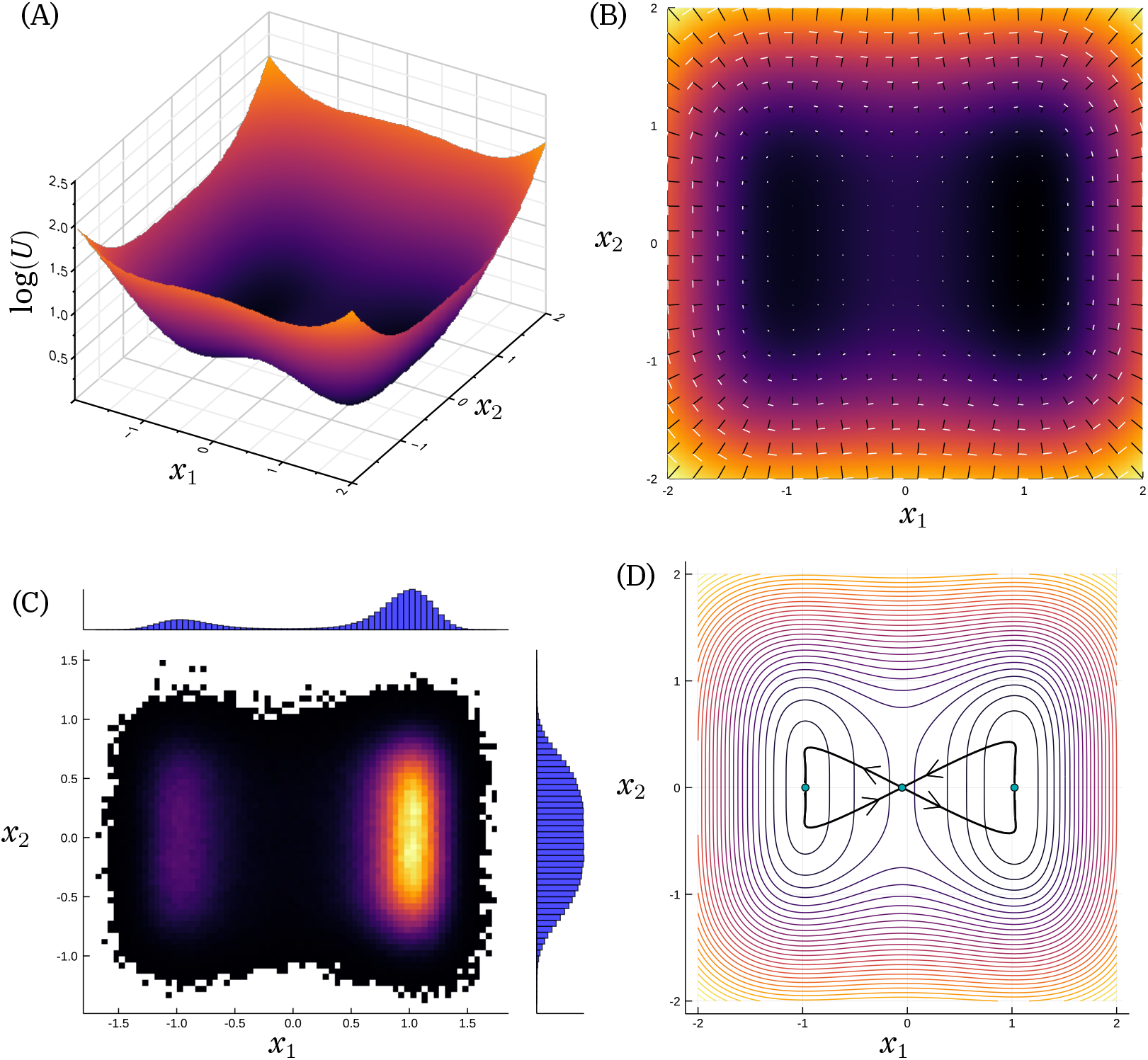
The two-dimensional SDE model: (A) surface plot of the potential field *U*; (B) the vector field decomposed into the gradient (black) and curl (white) components; (C) the two-dimensional histogram generated from a long-run simulation of the system; (D) The minimum action paths (MAP) between stationary points *X_i_*. On exit from an attractor the MAP follows a path *X* =*∇U*+*f_U_*, before following a “free fall” path back towards a stationary point.

### 1.2 Static landscapes and the transition state

The simple model discussed above consists of a fixed potential *U*, parametrized by the vector *θ* = [*λ*, *α*, *β*]. This potential defines a *static* landscape since the potential itself remains constant with time. Given such a static landscape, we can define a transition state (TS) for the system based upon the most probable transition paths between the two stable fixed points. In the limit of small noise, the transition paths follow a particular route, that of *least action* (Freidlin et al., 2012). These paths are known as the minimum action paths (MAP), and for a system in which the gradient and curl components are orthogonal, Freidlin and Wentzell further demonstrated that they follow a particular route: *X*^˙^ = *∇U* + *f_U_*. These paths are displayed in Figure 1(D).

The transition paths in both directions clearly pass through the unstable fixed point at *X* = (0, 0). This may therefore be identified as the transition state *X_t_*, and represents a location in state space that, with high probability, the system will pass through when transitioning between the two attractors of the system. In the context of stem-cell differentiation, such a state might represent a particular transcriptional configuration through which all cells transition when changing between a given two phenotypes. It is therefore important to consider which properties of the system determine how long is spent near this state, since this has implications for our ability to locate TSs, or map the path through gene expression space between cell fates from experimental single-cell data (Moignard et al., 2015; Stumpf et al., 2017; Moris et al., 2016)

In a dynamical systems context, the linear stability of fixed point such as *X_t_* can be determined from the Jacobian at this point. Here, we examine the eigenvalues of the Jacobian, which provide information regarding the local stability and typical trajectories following a small perturbation (Kwon et al., 2005). Eigenvalues with negative real part imply trajectories that decay towards the fixed point, while those with positive real part imply exponentially growing trajectories. In general, the real parts may be associated with the local landscape while the imaginary parts describe the locally orthogonal curl component.

For our example defined by equation 4, the positive real eigenvalue may be shown to be determined by the parameter *α*. We therefore simulate the system with two different values of *α*, while keeping the ratio *α*/*λ* constant. Example trajectories from such simulations are displayed in Figure 2.

**Figure 2:**
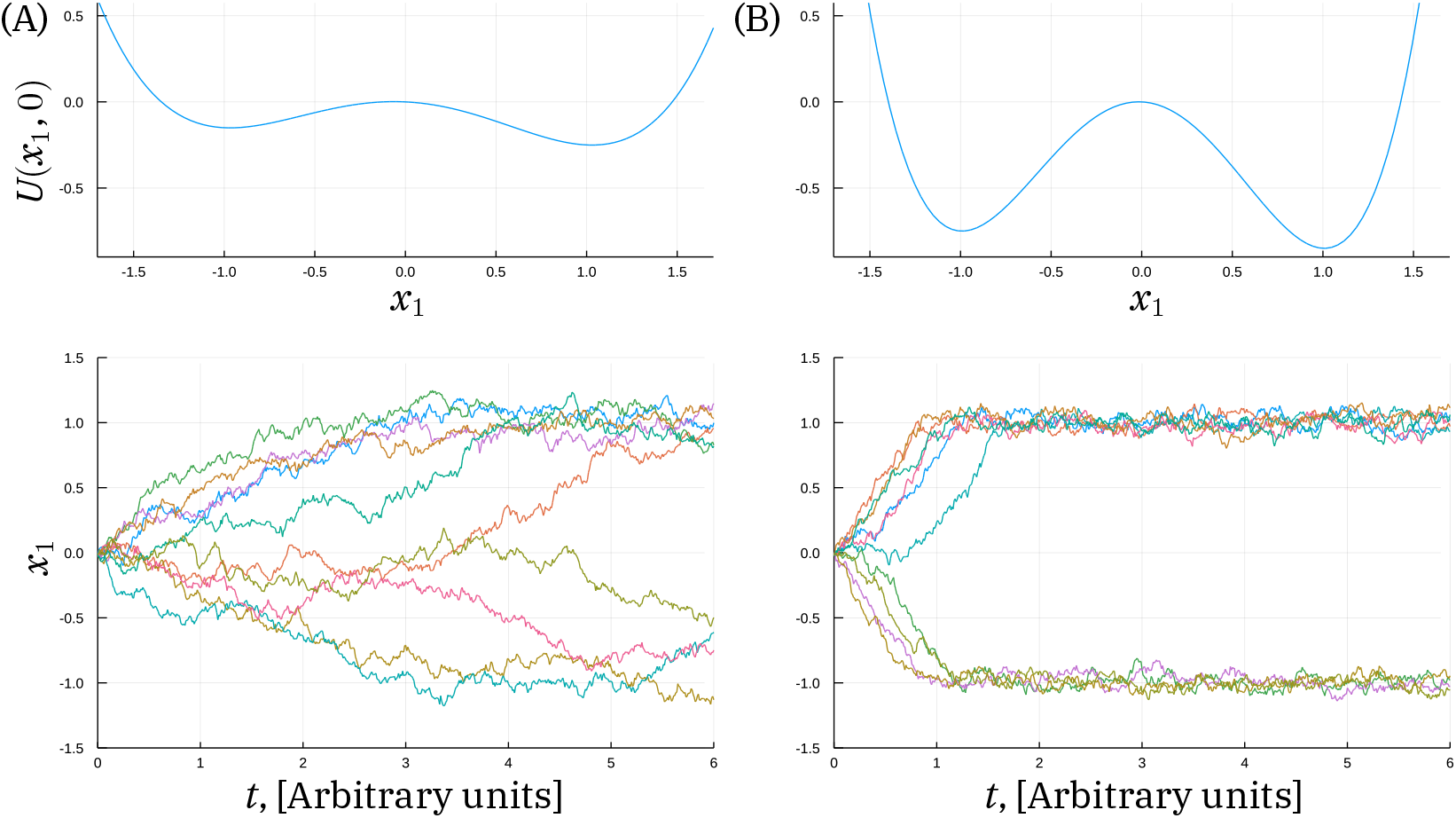
Slice through the potential and example trajectories away from the transition state under different model parameters: (A) small positive eigenvalue; (B) large positive eigenvalue. For a larger positive eigenvalue the wells of the system are deeper and the curvature is greater, leading to more rapid movement away from the transition state.

The trajectories shown in Figure 2(A), while all starting from the same initial conditions display high variability due to the stochastic nature of the system. Paths diverge towards one of the two stable equilibria for which *x*_1_ ≈ ± 1. The rate at which this divergence occurs varies considerably, although most seem to have undergone this divergence after 6 time units. By contrast, the trajectories in Figure 2(B) diverge towards the stable states in much less time, although the relative variability is similar. This may be attributed to the larger positive eigenvalue which governs the time-scale of the process. This is in turn related to the shape of the landscape as also shown in Figure 2: a larger positive eigenvalue leads both to a higher barrier in the potential between the two stable states as well as steeper roll off. The time spent at the transition state must reflect the curvature of the landscape, which in turn is quantified by the eigenvalues of the Jacobian.

### 1.3 Transitory landscapes: the competing definition for a transition state

While the system described above consists of a static landscape, a modified system might be formed by having parameters change over time. Here *β*alters the landscape, shifting which stable state is preferred. A landscape where parameters change may be referred to as *transitory*, as the shape of the potential is time-varying. In the context of epigenetic landscapes, the transitory interpretation gives an alternative viewpoint somewhat contrary to Waddington’s original, yet not without precedent in the literature (Wang et al., 2011; Rabajante and Babierra, 2015).

By varying *β* over a suitable range, the system changes from one in which there is a single stable equilibrium at positive *x*_1_ to the bistable system displayed in Figure 1, to one with a single stable equilibrium at negative *x*_1_.

We demonstrate the concept of a transitory landscape by generating an ensemble of simulations in which *β* varies linearly over time from −0.5 to 0.5. Figure 3(A) shows the variance across this ensemble of simulations for the state variables *x*_1_, *x*_2_. While the variance of *x*_2_ remains constant, the variance of *x*_1_ can be seen to begin and end at very low values but transition through a time-period of high variance. This high variance is associated with the bistability of the system that is observed for small absolute values of the parameter *β*, as displayed in the probability distribution for which *β* = 0.0 in Figure 3(B).

**Figure 3:**
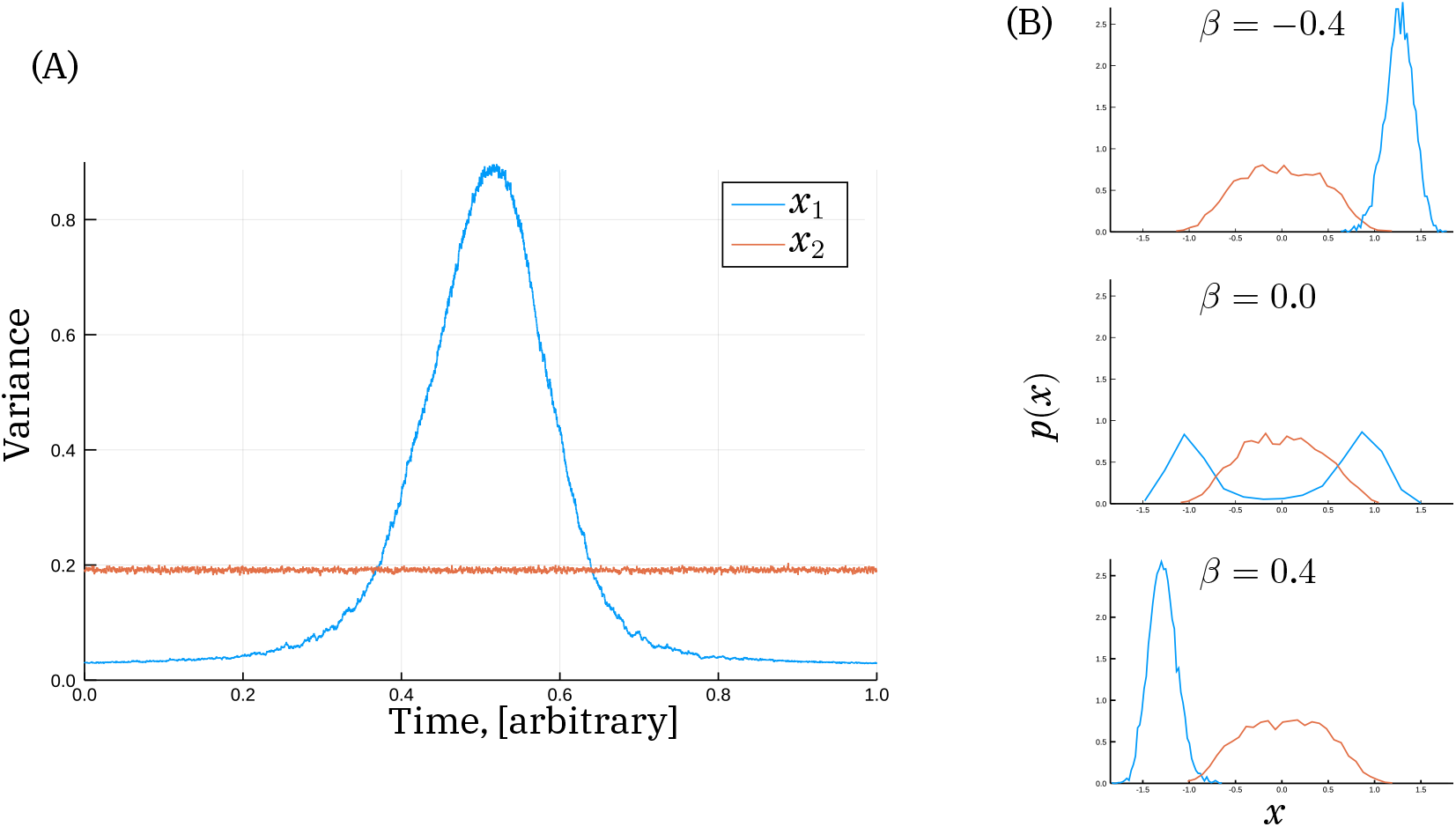
The transition state in a transitory landscape: (A) The variance of the two states across 5000 simulations in which βis linearly varied with time; (B) Probability distributions of the two states at three different values of β. The high variance “transition state” corresponds to intermediate values of β in which significant bistability generates large heterogeneity.

The time-period of high variance displayed in Figure 2(A) is analogous to the transition state concept posited by Moris et al. (2016). In the context of stem cell development, high variance across an ensemble of simulations is equivalent to high heterogeneity across a population of cells at the same developmental stage.

It is also worth noting at this point that the choice between a static or transitory landscape is really an issue of model complexity: as an alternative we can consider *β* to be an additional state rather than a model parameter. The system would therefore be three-dimensional with some additional dynamics governing how this third state *x*_3_ = *β* evolves with time. Both models will be able to capture the same behaviour but would differ in the way they described it: the landscape of the former being two-dimensional and time-varying while that of the latter being static and three-dimensional.

## 2 The Transition States and Landscapes in a Developmental Model

With the insights derived from these illustrative models, we continue our analysis with a model that incorporates some of the dynamic relationships involved in stem cell differentiation (Chickarmane et al., 2012). The model focuses on four key players: Nanog (*N*), the complex Oct4-Sox2 (*O*), Fgf4 (*F*), and Gata6, (*G*) a typical differentiation marker. A schematic representation of their network is shown in Figure 4(A); all arrows stand for non-linear interactions that account for implicitly modelled species and processes, like the formation of the Oct4-Sox2 complex, or Nanog dimerization. In addition, Leukaemia inhibitory factor (LIF), is also included in the model: the concentration of LIF (*L*) is treated as an external control, through which the environment of the cells is modified. Full details of the model are given in the methods section, while further analysis may be found in Lakatos (2017).

**Figure 4:**
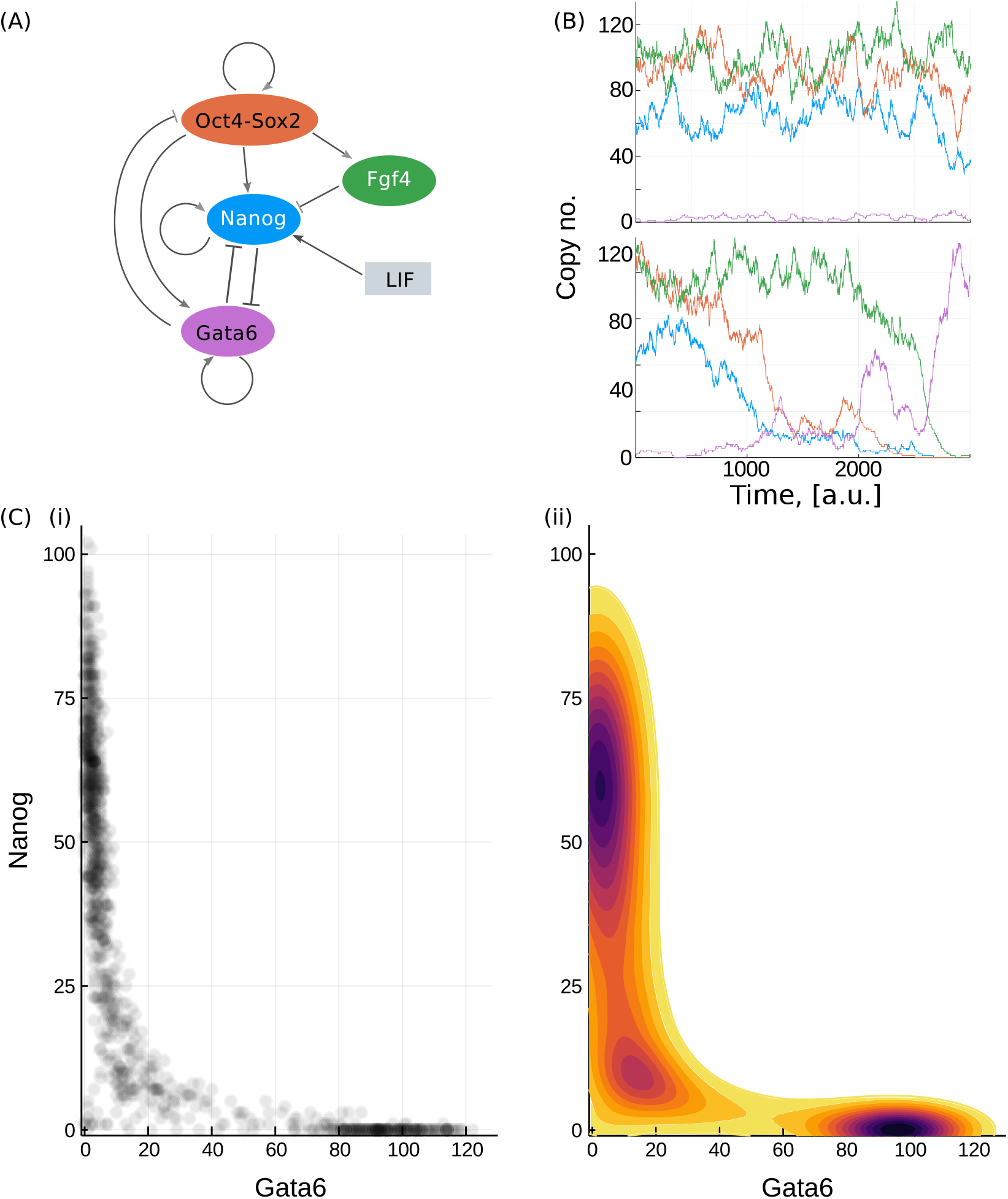
Potential landscape of the stem cell differentiation model. (A) Schematics of the stem cell differentiation model. (B) Simulated example of a cell that stays pluripotent (upper axes) and a cell undergoing differentiation (lower axes). The abundance of each species is indicated by a line coloured according to the filled ovals in (A). (C) Probability distribution and landscape of the system with L = 50. (i) Scatter plot of the whole population on the Gata6-Nanog plane. (ii) The landscape computed as the negative logarithm of the probability distribution.

### 2.1 The probabilistic landscape and system behaviour

In this model, high expression of Nanog and Gata6 indicate stem cell and differentiated phenotypes, respectively. Since there is mutual inhibition between these two molecules, a state (reached under deterministic dynamics) where both are simultaneously high or low is unlikely. Figure 4(B) shows example simulations with *L* = 50 and intermediate initial expression levels, under which conditions many cells stay pluripotent (i), but some stochastically shift into a differentiated state (ii). In general, simulations initiated with high values of Nanog, Fgf4 and Oct4-Sox2 will fluctuate around these high values, before ultimately switching to a configuration with high Gata6 and low expression levels of the other molecules. The time at which this switch occurs is highly variable, but the switch itself is seen to be irreversible, indicating that the differentiated state is in some sense more stable.

In order to gain further insight into the behaviour of the system we compute the probabilistic landscape for *L* = 50. This computation is achieved by performing a large number of simulations over a fixed time-period, from which the final expression levels is sampled. The landscape is then evaluated as minus the logarithm of this empirical distribution and is shown in Figure 4(C).

The potential landscape in Figure 4(C) displays a narrow valley which provides the most likely connection between the differentiated and pluripotent states. To compare the final potential landscape with paths taken by cells, let cells differentiate *in silico*. First, we start the cell population in a high LIF medium, with initial values from the *N* ∈ [60, 100], *G* ∈ [0, 16] region and *L* = 200. We then change the external conditions, *L*: 200 → 0, and simulate the differentiation paths. As demonstrated by randomly selected trajectories in Figure 5(A), the dynamical paths show good correspondence with the previously derived static landscape: cells traverse through the narrow valley between the stem cell and differentiated states.

**Figure 5:**
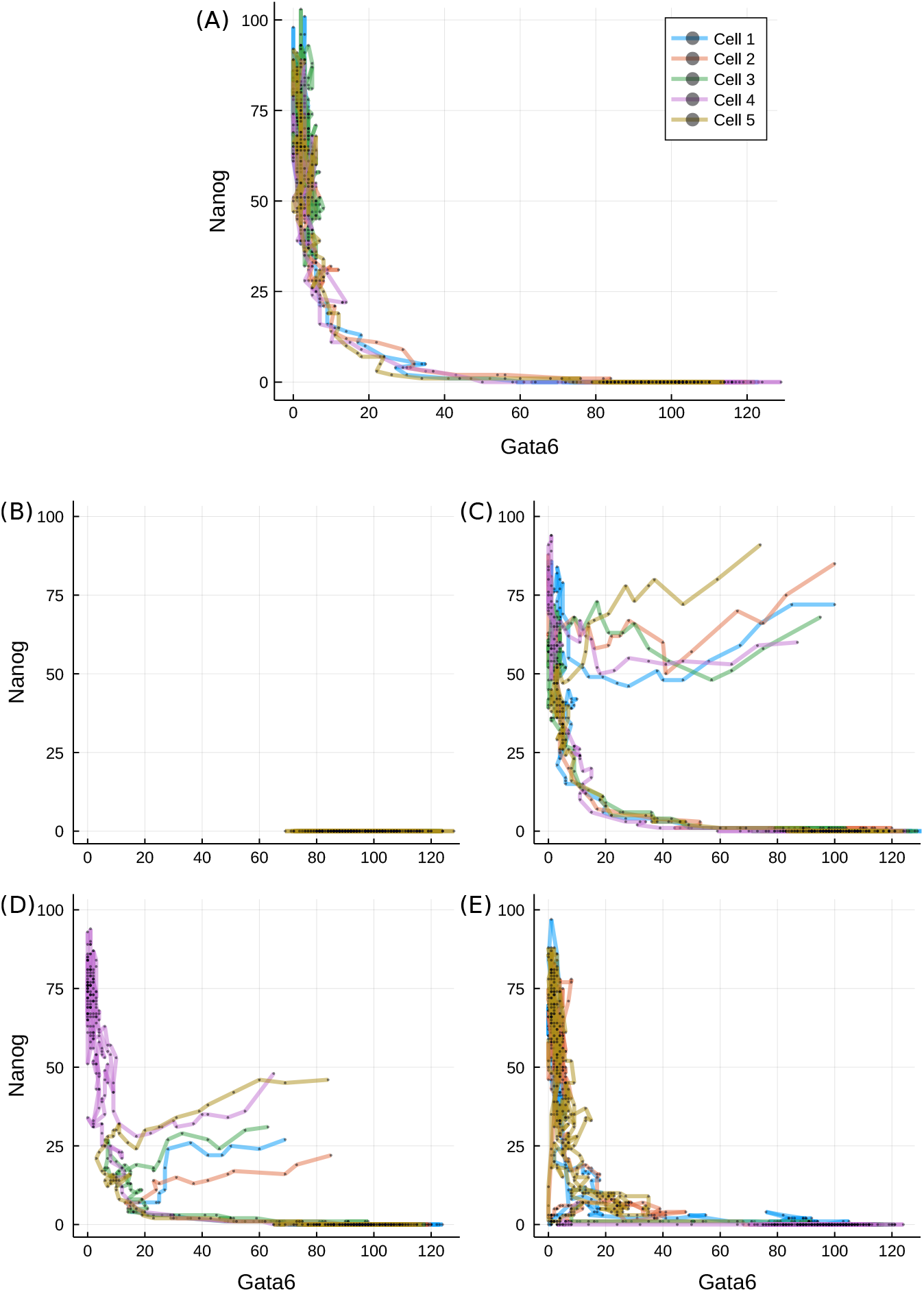
Transition paths during differentiation. (A) Trajectories of five randomly selected stem cells after reduction in LIF concentration (L = 0), plotted on the Gata6-Nanog phase space. (B) Cells started from the differentiated state and put into L = 200. (C-D) Cells started from the unexplored region of high Gata6 and high Nanog (C) or low Nanog (D) levels for L = 0. (E) Trajectories started from the transition state under L = 50. Different line colours denote different cells, as indicated in the legend.

Most of the time along typical cell paths is spent in either of the two valleys: cells linger in the stem cell valley for varying amount of time, but as their Nanog level decreases below a critical threshold, they quickly transit to the differentiated cell valley. Some trajectories do not exit the self-renewing state (such as cell 3 on panel (A)) over the simulated time-course, accounting for the less populated but still prevailing stem cell population in Figure 4(C). We also test the system in a reversed differentiation scenario: cells already in the differentiated valley are simulated under high LIF conditions (*L* = 200) and tracked. The trajectories confirm that once differentiation is achieved the cells cannot regain pluripotency, as consistent with an irreversible switch-like process.

As is clear from the derived landscape, the majority of the Gata6-Nanog phase space lies on a high-potential plateau that remains unexplored by the cells, displayed as the large white region of Figure 4(C). Therefore, we have to artificially introduce conditions that allow the mapping of cell dynamics on the value range of this plateau. We first uniformly sample cells from an uncovered volume of four-dimensional space, for example {*N*, *O*, *G*} ∈ [60, 100] and *F* ∈ [10, 50]. Next we follow the same experimental procedure as before and induce differentiation by setting *L* to 0. Surprisingly, trajectories from this region quickly converge into the Nanog-high valley and then follow the paths observed on a stem cell starting population, as displayed in Figure 5(C). On the other hand, initial values sampled from another region of the high-potential plateau ({*O*, *F*, *G*} ∈ [60, 100]; *N* ∈ [20, 50]) lead the majority of cells towards the differentiation basin, but do so via the TS. Nevertheless, there is a significant fraction of cells, solely with initial Nanog abundance over 40, that go towards and linger in the stem cell state.

Interestingly, as illustrated in Figure 5(D), cells undergoing differentiation do not follow a direct route. They first visit a state characterised by simultaneously low Gata6 and Nanog expression, then follow the connecting valley to a differentiated phenotype. Such a state is reminiscent of the static transition state discussed above, while the phenomenon is consistent for a wide range of initial conditions, suggesting that any cell – unless it has already differentiated – has to go through the transition state. Cells that reside in the transition state, on the other hand, have the ability to progress towards either states, as seen in Figure 5(E). This is in agreement with other models reporting a distinguished transition state along the way of cell specialisation that has the potential for any cell fate (Martinez Arias and Hayward, 2006; Muñoz Descalzo and Martinez Arias, 2012; Li and Wang, 2013).

### 2.2 Optimal paths in the static landscape

The indirect transition paths observed to be taken by the cells are indicative of a curl component in the dynamics. This finding does not contradict the landscape of Figure 4(C) since such components are not captured in the potential, but is illustrative of the ubiquity of such dynamics.

One implication arising from the presence of curl is that the optimal paths in the forward and reverse directions will not be identical, as was seen for the toy model in Figure 1(D). For this system the optimal, or minimum action, path in the forward direction may in principle be inferred from simulations, since this path is also the most probable. However in practice the high variability between simulations makes the precise path difficult to observe. In the reverse direction, simulations may never yield a path, and the hypothetical MAP must instead be evaluated via an optimisation approach, as detailed in Perez-Carrasco et al. (2016). The MAPs in each direction are displayed in Figure 6.

As is clear from Figure 6(A), the paths in the forward and reverse directions are different, although both traverse the valley-like region of the Gata6-Nanog landscape shown in figure 4(D). While the paths are similar, it is important to note that in the four-dimensional space of this model, they never meet except at the start and end points. However both paths come close to each other near to the static transition state at low values of both Gata6 and Nanog. While the reverse path travels to and from the transition state with more-or-less direct trajectories, the forward path is seen to loop around in a convoluted manner. Such behaviour is indicative of curl dynamics and is consistent with the complex eigenvalues of the system, as will be discussed below. The differences between the forward and backward paths may be seen more clearly in the time-series. In the forward direction, the MAP begins with a slight transient increase in Fgf4 accompanying a gradual fall in Nanog and Oct4. Gata6 rises slowly at first before rising much more rapidly once Nanog is below a critical level. The mutual inhibition between Nanog and Gata6 leads to a similar effect in the reverse direction: Gata6 must fall rapidly, and only once it is below a fairly low level (≈ 10) does the abundance of the other species begin to rise (Figure 6(C)).

**Figure 6:**
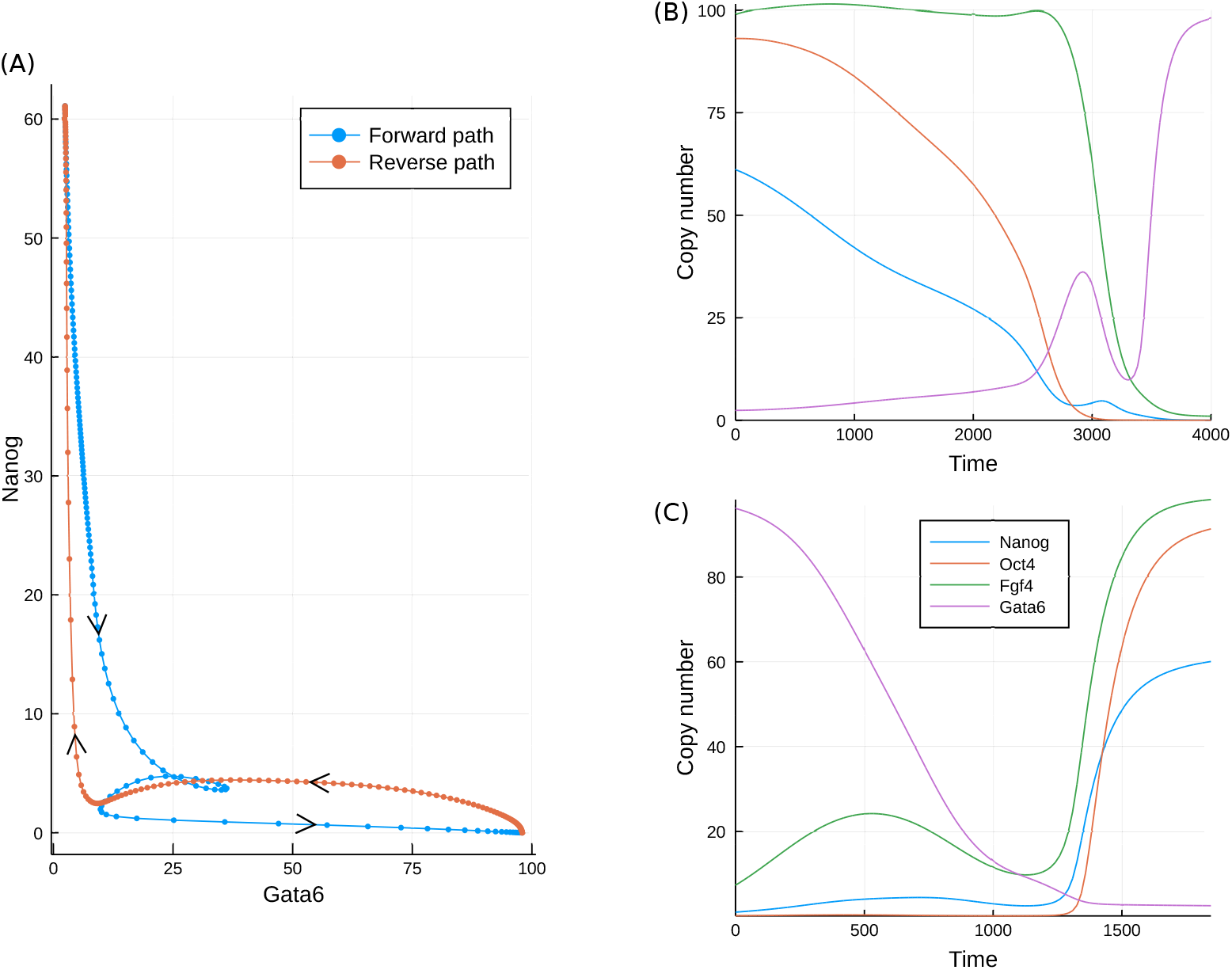
The minimum action paths for the developmental model in both the forward and reverse directions. A comparison of the paths plotted in the Gata6, Nanog plane; (B) Time series for each of the molecular species in the forward direction; (C) Time series for each of the molecular species in the reverse direction.

The key observation from the MAPs is that because of the curl component to the dynamics, the most probable paths and static transition states are different for each direction. In some cases the paths meet at a particular state such as a saddle point in the landscape (Lv et al., 2014). However in general this will not be the case, even for relatively simple low-dimensional systems (Wang et al., 2010b, 2011; Li et al., 2016). Thus, if we wish to gain insight into the easiest routes for the reprogramming of cells back to pluripotency, observation of the typical forward paths will likely not be sufficient.

### 2.3 Transitory landscapes in a time-varying model

The behaviour illustrated above suggests the existence of a transition state in the static landscape, similar to the saddle point in the toy model discussed above. However, the transition state may also be defined for a transitory landscape, which in this case may be achieved by varying the level of LIF, quantified by the parameter *L*.

Before proceeding to simulations of a transitory system, it is first insightful to examine analytically how the fixed points (*X_i_*) of the system vary with *L*. This is achieved by finding the expression levels at which the rate equations are equal to zero. Three such fixed points are displayed in Figure 7(A) for varying *L*. The three identified fixed points may be associated with each of the stem-cell, transition and differentiated states of the system. Of these three states it is only the stem-cell state that is strongly influenced, moving to higher values of Nanog as *L* is increased.

**Figure 7:**
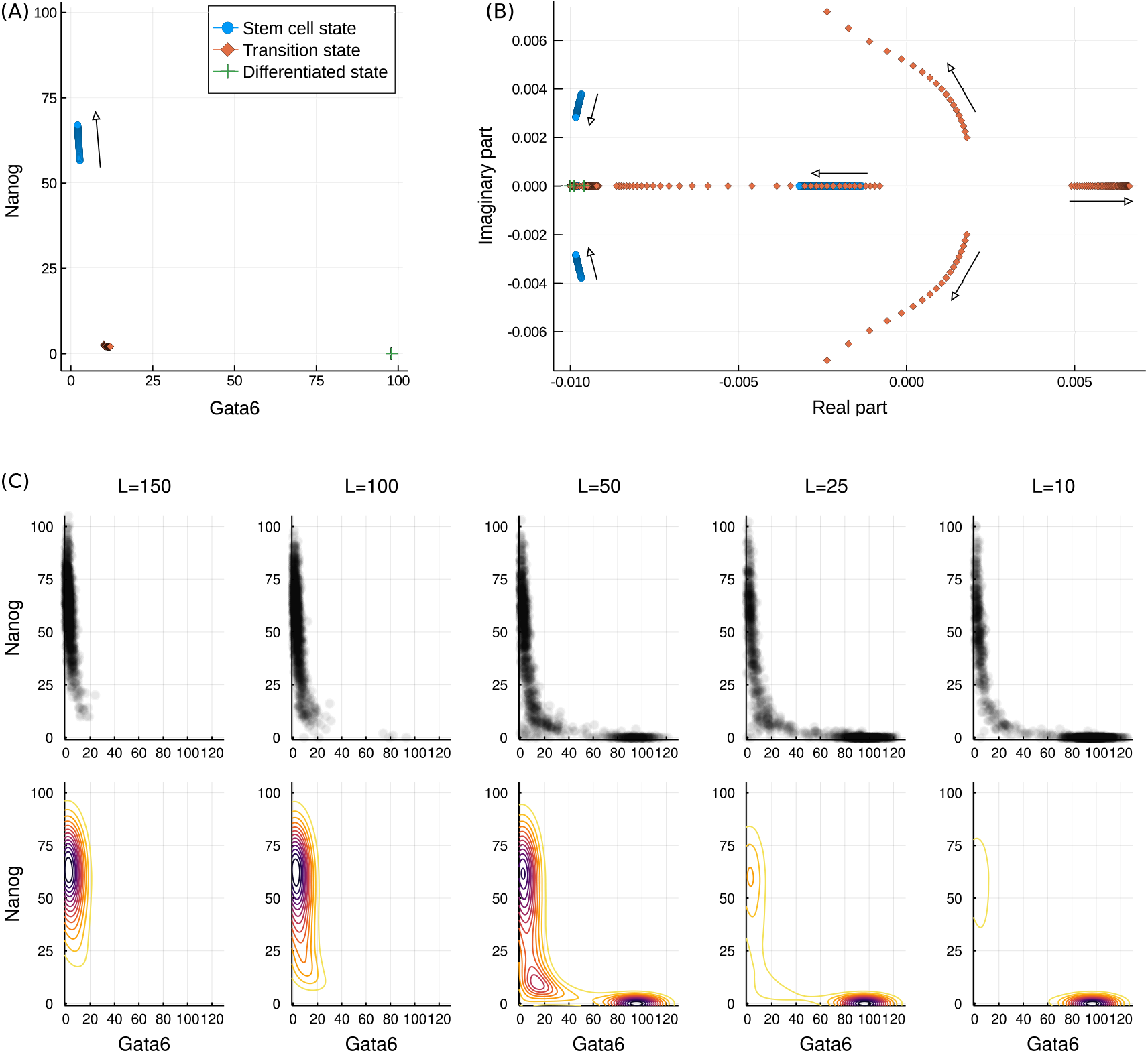
Varying behaviour of the system with LIF concentration, L. (A) Three of the fixed points of the model with increasing L. The differentiated and transition state remain unchanged while the stem cell state moves to higher Nanog values. (B) The eigenvalues of the Jacobian at each of the three fixed points as L increases. The stem cell state becomes locally more stable, as indicated by the increasingly negative eigenvalues. The transition state becomes increasingly unstable, thereby making transition to this state less probable. The differentiated state is unaffected, but is seen to be locally more stable than the stem cell state. Complex eigenvalues are associated with rotational “curl” dynamics. (C) The changing probabilistic landscape. Upper plots show a sample of the sampled points while lower plots show the landscapes. As L decreases the valley at the differentiated state becomes deeper while that at the stem cell state becomes shallower. At intermediate values the landscape is flatter around the transition state.

**Figure 8:**
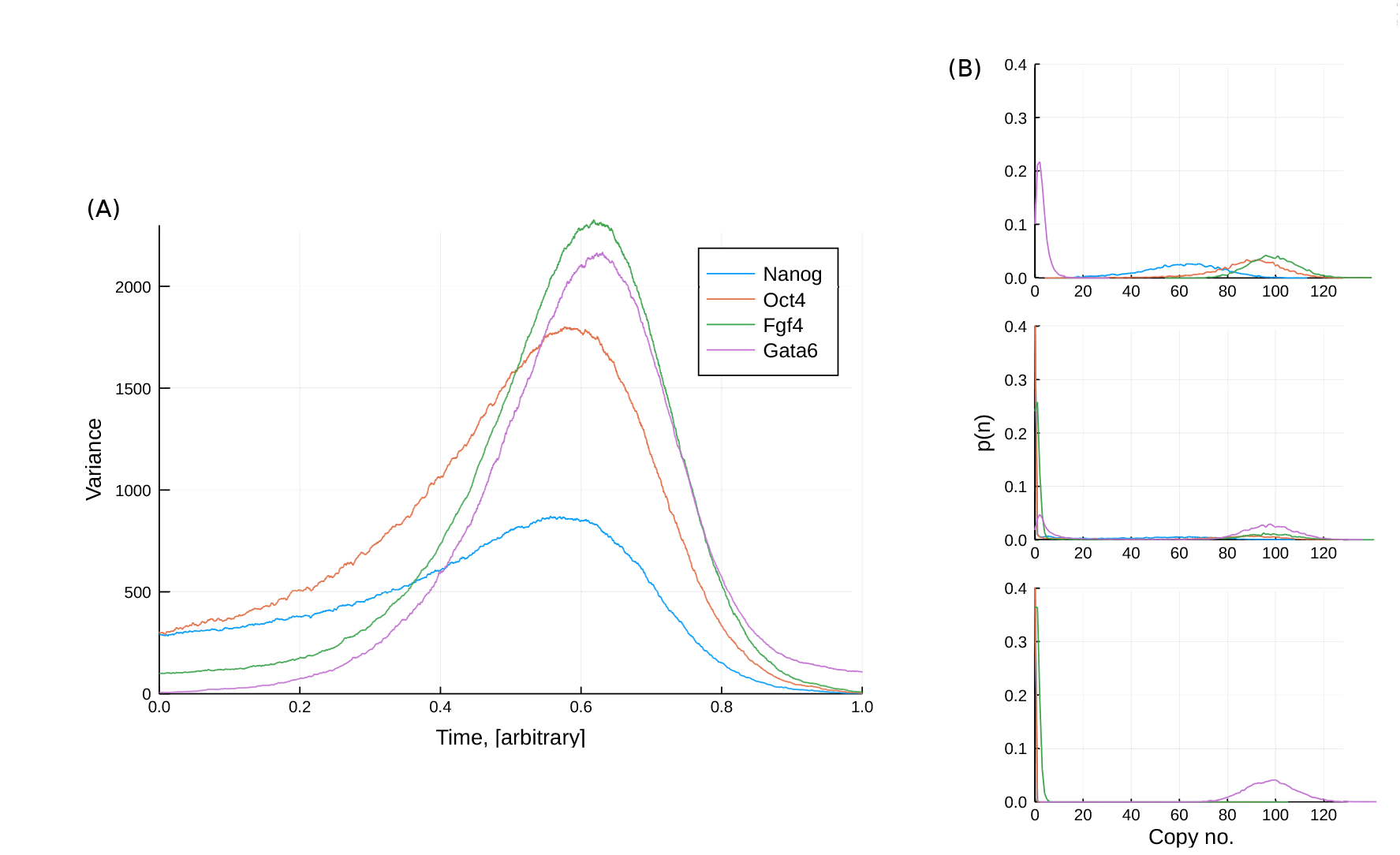
The transition state in a transitory landscape: (A) The variance of the transcription levels across 10000 simulations in which L is linearly decreased with time; (B) Probability distributions of the transcription levels at three different values of L. The high variance “transition state” corresponds to intermediate values of L at which transitions from the stem cell to differentiated state can occur.

In addition to the location of the fixed points we may also again examine the eigenvalues of the Jacobian at each of them. These eigenvalues, over a range of LIF concentrations, are displayed in Figure 7(B) as a scatter plot in the complex plane. The eigenvalues for the stem cell and differentiated state can be seen to reside entirely in the left half plane, a characteristic indicative of the linear stability of these states. In contrast the transition state is seen to have one eigenvalue with positive real part (and is hence unstable). Furthermore, the magnitude of this eigenvalue increases with increasing *L*. This implies both that the system will leave this state more rapidly if it passes through it, but also that arriving at this state is less probable, consistent with the observation that high *L* inhibits (initiation of) differentiation.

The eigenvalues of the stem-cell state also vary with LIF concentration, in this case transitioning to complex values with increasingly negative real part. This implies increased stability of this system and, together with the variation of the transition state eigenvalues, is consistent with the behaviour of the system under increasing *L*: increased stability of the stem cell state.

The eigenvalues of the differentiated state do not vary with *L*, but all the values are seen to be more negative than some of those of the transition and stem cell states. This is consistent with the greater stability of the differentiated state and the tendency for the system to remain there once it has arrived. Furthermore, these properties of the different states may be related to the shape of the probabilistic landscapes shown in Figure 7(C). As *L* varies the landscape is seen to transition from one with a single valley around the stem-cell state to one with two valleys of varying depth. Further to this, for the case of *L* = 50 the differentiated state is surrounded by more tightly spaced contours than is the stem-cell state, and hence higher local curvature; this is consistent with the relative size of the eigenvalues.

Given the observed variation of the system behaviour, we run simulations in which the parameter *L* is varied linearly over time. As for the toy model in section 1, we examine the variance across a large ensemble of simulations (10,000). This is displayed in figure 8. The ensemble starts out with relatively low variance before displaying a period of high variance at intermediate times. In the context of real experimental data, such high variance would be observed as large heterogeneity across the sampled cells. This heterogeneity occurs because for this particular model, there is a distribution of times at which the stochastic switch is made from the stem-cell to differentiated state. Therefore even though all cells follow a similar path through gene expression space with very similar start and end points, there is high variability between cells over a particular period of time.

## 3 Discussion

Models describing stem-cell differentiation are plentiful (Herberg and Roeder, 2015). In this work we have examined one such model, that of Chickarmane et al. (2012), from a landscape perspective. Starting our analysis from an illustrative model we see that the landscape is strongly linked to the probability distribution over the state space, and gives an intuitive description of the dynamics. In the stem-cell differentiation model we are able to obtain a landscape that describes the typical system behaviour. Moreover, the landscape may be linked to the dynamics in a quantitative manner via the fixed points *X_i_* and the eigenvalues of the linearised dynamics at these locations: the fixed points occur at minima, maxima or saddle points in the landscape while the eigenvalues describe the local curvature.

While the potential landscape describes the tendency of the system to move towards particular stable states, it cannot fully capture all of the system’s dynamics. A non-gradient contribution to the dynamics is also generally present, known as the curl. A curl component is thought to be indicative of non-equilibrium systems, or those which are not closed to external influences (Wang, 2015). This is because closed systems always have an available energy such as the Gibbs free energy which may be quantified and is directly related to the dynamics. For a developing stem cell, there is a constant exchange of both energy and nutrients via the diffusion of heat and mass through the membranes of the cell. Such a system is therefore clearly not closed in a thermodynamic sense, and non-equilibrium effects are bound to be present.

An alternative viewpoint on the presence of curl, is that a curl-free or purely gradientbased system requires complete symmetry in the interactions between all state variables (Weinreb et al., 2017). In the context of gene regulatory networks, a necessary condition is that the influence of gene *i* on the expression of gene *j* is exactly the same as in the opposite direction. However even when this is the case, only in cases where these influences are of a particular functional form will the necessary symmetry be present.

The precise requirement is that 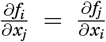, since both are equal to 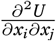. For systems such as a symmetric toggle switch in which there is mutual repression between two gene products, the standard Hill function form of the interaction does not lead to this symmetry. Therefore in general, with the exception cases in which the regulatory equations are specially chosen (Olariu et al., 2017), we may expect a curl component to the dynamics of the system.

Although the potential landscape cannot describe all of the dynamics of cell differentiation, it can elucidate some pertinent features of the process. One such feature is the static transition state. Given a static landscape description for a developing cell, we may define transition states as the saddles in the landscape. The behaviour of the system at these transition states is linked to the eigenvalues of the linearised dynamics, evaluated at these states. Such saddle points will always have eigenvalues with a positive real part, the magnitude of which is linked to the speed at which the transition states may be traversed. In the case that the transition speed is quick, the transition states may rarely be observed and measured trajectories will appear discontinuous.

The static transition states may also be inferred without recourse to the landscape, utilising the concept of the minimum action path. Such paths have been evaluated for similar developmental models before (Zhang and Wolynes, 2014), and are in general agreement with those found here. The MAPs describe the most probable routes through state space between any two fixed points. For purely gradient-based systems these routes simply follow that of steepest climb/descent and are therefore identical in both directions between any two points. For two-dimensional systems with an additional curl component such as the toy model examined above, the paths differ between the two directions but will meet at the saddle point between adjacent attractors. Such a saddle point may therefore be termed a static transition state. For higher dimensional systems such as the developmental model, the paths are again different but still come close together near to one of the fixed points. The transcriptional region around this static transition state is therefore of significance for both differentiation and reprogramming.

An alternative to the static landscape description of the system is one of a transitory landscape, in which the potential changes under the influence of an external input or time-varying parameter. In this framework we may therefore consider an alternative definition for the transition state as those periods during development in which large heterogeneity is observed. Such heterogeneity may arise from two effects, firstly from the broad distribution of times for the transition from one fixed point to another, and secondly from a temporary flattening of the landscape under the influence of the changing conditions. In the models discussed here, both effects are present. The variation in switching times is typical of any stochastic system in which the escape from an attractor occurs due to random perturbations (Gammaitoni et al., 1998) and is therefore an inherent feature of many processes governed by a gene regulatory network. The temporary flattening of the landscape is dependent on the particular nature of the interactions between genes, but there is increasing experimental evidence for such a phenomenon (Martinez Arias and Brickman, 2011).

When forming any gene regulatory model, the choice between a static and transitory landscape may often be an issue of model complexity. For a gene regulatory network this choice is akin to that between modelling all of the relevant network interactions, or simply taking a subset of the network and treating the influence of other genes as (time-varying) parameters in the model. In the particular application of stem-cell development there is evidence that a relatively limited number of genes may be sufficient to describe typical behaviour, although external inputs are still required (Dunn et al., 2014). Such a system may therefore be described by a transitory landscape.

An important final consideration arising from either landscape interpretation is that the presence of curl places limits on the identifiability of the dynamics. Most experimental data consists of purely static snapshots, from which instantaneous distributions can be obtained. Methods such as scRNA-seq only allow measurements of transcriptional profiles at single instances, and cannot provide the temporal variation of any individual cell. While this will enable measurement of the typical distribution of cell states, it will not readily allow measurement of certain characteristics of the cell paths, as discussed in Weinreb et al. (2017). The full identification of regulatory dynamics may therefore additionally require some measure of derivatives, as may be afforded by future experimental and analytical techniques (La Manno et al., 2017); this will still leave considerable challenges to identifying model structures (Babtie et al., 2014).

## 4 Methods and Materials

### 4.1 Two-Dimensional Stochastic Model

The two-dimensional toy model is chosen to exemplify some key properties of nonlinear stochastic dynamical systems. It is a two-dimensional extension of a well studied one-dimensional bistable system Gammaitoni et al. (1998), known to exhibit a wide range of phenomena. The governing equations are,

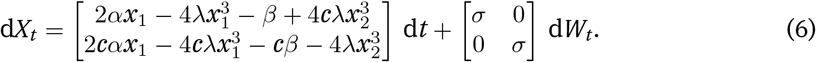

The parameter values used in the model were

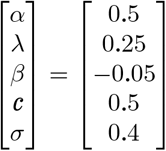

### 4.2 The Developmental Model

The developmental model consists of four key players: Nanog (*N*) the complex Oct4-Sox2 (*O*), Fgf4 (*F*) and Gata6 (*G*). We additionally include the typical cell media LIF (*L*), which is a controllable parameter. Applying the quasi-equilibrium assumption, we treat all modifying interactions as factors in the same birth process, with their associated rates acting as a weighting on the net production rate. A first order degradation reaction parametrised by *k_d_* is also included for each molecular species, leading to eight reactions with the following propensities:

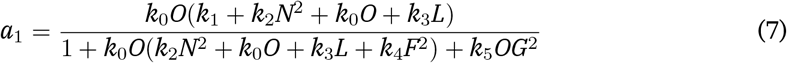

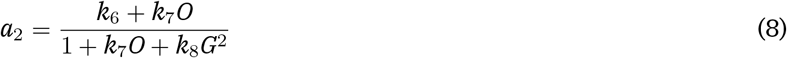

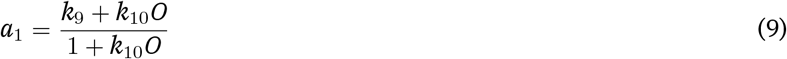

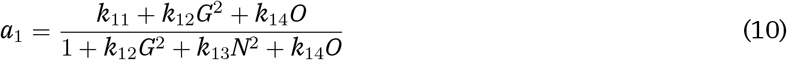

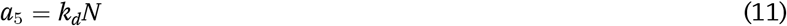

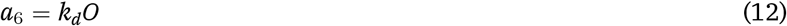

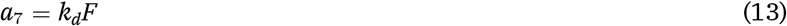

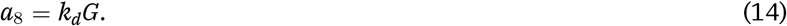

The evolution of the system is then described by the combination of these eight reactions and a stoichiometry matrix *S*, that defines the integer changes to the copy number of each species. This is given as,

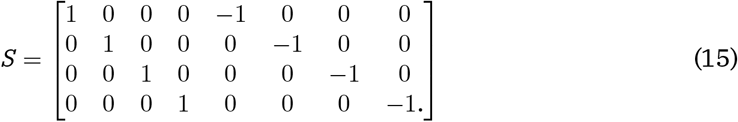

The parameter values used in the model were

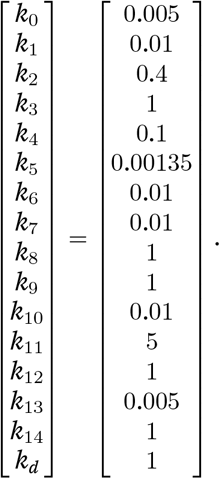

### 4.3 Numerical and analytical methods

Simulations of the developmental model were performed using the Gillespie algorithm Gillespie (1976), as is standard for chemical reaction systems. For the analysis of the fixed points, linearised dynamics and minimum action paths, the system was transformed into the approximate SDE format. Defining the vector *X* = [*N*, *O*, *F*, *G*]^⊤^, and the reaction vector *a* = [*a_i_*]^⊤^, the SDE approximation is given as

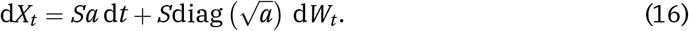

Here diag(*v*) stands for the ℝ^*N*×*N*^ matrix with the elements of vector *v* in its diagonal and zeros elsewhere.

The deterministic fixed points of the system may be obtained by solving the equation,

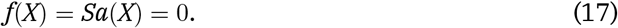

Due to the nonlinear nature of the equations, analytical solutions are difficult to obtain. Solutions were therefore found numerically using the `DifferentialEquations.jl` package in Julia (Rackauckas and Nie, 2017). Analysis of the Jacobians was performed symbolically using the `SymPy.jl` package.

The minimum action paths were computed via the optimisation approach detailed in the supplementary information of Perez-Carrasco et al. (2016). The MAP are those paths through the state space that minimise the Friedlin-Wentzell action functional (Freidlin et al., 2012).

## Acknowledgements

We are grateful to members of the theoretical systems biology group for helpful discussions.

